# Persistent thermal input controls steering behavior in *Caenorhabditis elegans*

**DOI:** 10.1101/2020.04.29.067876

**Authors:** Muneki Ikeda, Hirotaka Matsumoto, Eduardo J. Izquierdo

## Abstract

Motile organisms actively detect environmental signals and migrate to a preferable environment. Especially, small-size animals convert subtle difference in sensory input into orientation behavioral output for directly steering toward a destination, but the neural mechanisms underlying steering behavior remain elusive. Here, we analyze a *C. elegans* thermotactic behavior in which a small number of neurons are shown to mediate steering toward a destination temperature. We construct a neuroanatomical model and use an evolutionary algorithm to find configurations of the model that reproduce empirical thermotactic behavior. We find that, in all the evolved models, steering rates are modulated by persistent thermal signals sensed through forward locomotion. Persistent temperature increment lessens steering rates resulting in straight movement of model worms, whereas temperature decrement enlarges steering rates resulting in curvy movement. This relationship between temperature change and steering rates reproduces the empirical thermotactic migration up thermal gradients and steering bias toward higher temperature. Further, spectrum decomposition of neural activities in model worms show that thermal signals are transmitted from a sensory neuron to motor neurons on the longer time scale than sinusoidal locomotion of *C. elegans*. Our results suggest that employments of persistent sensory signals enable small-size animals to steer toward a destination in natural environment with variable, noisy, and subtle cues.

**Author summary:** A free-living nematode *Caenorhabditis elegans* memorizes an environmental temperature and steers toward the remembered temperature on a thermal gradient. How does the *C. elegans* brain, consisting of 302 neurons, achieve this thermotactic steering behavior? Here, we address this question through neuroanatomical modeling and simulation analyses. We find that persistent thermal input modulates steering rates of model worms; worms run straight when they move up to a destination temperature, whereas run crookedly when move away from the destination. As a result, worms steer toward the destination temperature as observed in experiments. Our analysis also shows that persistent thermal signals are transmitted from a thermosensory neuron to dorsal and ventral neck motor neurons, regulating the balance of dorsoventral muscle contractions of model worms and generating steering behavior. This study indicates that *C. elegans* can steer toward a destination temperature without processing acute thermal input that informs to which direction it should steer. Such indirect mechanism of steering behavior is potentially employed in other motile organisms.

## Introduction

Animals sense environmental signals and navigate to a preferable environment [1,2]. Even when the distribution of signal is not known in advance, animals move around and detect a spatial signal gradient, enabling adjustments of moving direction for navigation. Such an active sampling of environmental signals is an essential component of the spatial navigation strategy for small-size animals, since it is difficult to detect a difference of signal intensity through multiple sensory organs placed on their tiny body [3,4].

With only a 1-mm-long body, the nematode *Caenorhabditis elegans* can sense and navigate gradients of gustatory, olfactory, and thermal signals [5–7]. During navigation, *C. elegans* makes gradual adjustments of its moving direction to steer upward/downward in a gradient [5,8,9]. Recent studies suggest a neural mechanism of steering behavior; worms could adjust the amplitude of head swings by sampling the difference of signal intensity through their own dorsoventral sinusoidal motion [10,11]. Supporting this hypothesis, oscillatory activity of premotor interneurons and motor neurons, which synchronizes with dorsoventral head swings, are modified upon the application of a favorable olfactory signal [12–14]. Further, optogenetic manipulation of a series of neurons only when the animal swings their head to dorsal or ventral side generates steering behavior to one direction [13,15,16]. However, during behavioral assays of freely moving animals, the difference of signal intensity sensed through head swings and resulting neural responses should be much more subtle. Especially in thermotaxis assays, temperature difference along one head swing is less than 0.01°C on a linear thermal gradient of 0.5°C/cm [7], which mimics natural thermal gradients in the upper few centimeters of soil [17]. Therefore, a novel mechanism potentially underlies steering behavior in such shallower signal gradients, albeit there had been no alternative hypotheses.

Here, we show that thermal inputs sensed not through head swings but through forward movement of *C. elegans* can generate steering behavior, leading worms to preferred temperature. To examine how the steering behavior is regulated during thermotaxis, we construct a neuroanatomically-grounded model with a set of neurons shown to mediate thermotactic steering behavior [9]. Thermal input sensed by the model worm was converted to the activity of a thermosensory neuron AFD through an empirical response property [18]; inter- and motor neurons were mathematically modeled as passive isopotential nodes with simple first order nonlinear dynamics [19,20]; and steering rates were assumed to be proportional to the difference in activities of dorsoventral neck motor neurons. The unknown electrophysiological parameters of the model, including the sign and strength of the connections, were optimized by running a large set of evolutionary searches [21] so as to reproduce the empirical thermotactic behavior [9]. We found that in all the ensemble models obtained through evolutionary searches, steering rates of model worms were modulated by the thermal input at the scale of forward movements rather than the scale of head swings. Temperature increment through forward movements lessened steering rates resulting in straight movement, whereas temperature decrement enlarged steering rates resulting in curvy movement. Our simulation analysis demonstrated that the observed relationship between temperature change and steering rates can reproduce the empirical steering bias and thermotactic migration. Further, spectrum decomposition of neural activities in model worms showed that the dynamics of temperature signal was transmitted from a sensory neuron to motor neurons on a longer time scale than head swings. Our results suggest that the signals sensed through forward movement allows worms to adjust their direction of movement such as to implement navigation to a preferable environment, without knowing the specific steering direction.

## Results

### Neuroanatomical models reproduce steering behavior during thermotaxis

*C. elegans* is known to navigate using a series of stereotyped movements: turns, reversals, and curves [5,8,22]. Worms bias their frequency of turns and reversals, thereby migrating indirectly to a destination; worms are also reported to bias their curving angle and direction of locomotion, thereby steering to a destination. Steering behavior is observed in chemotaxis [5,8,23], electrotaxis [24], and thermotaxis [9,25]. A recent study identified neural circuits for thermotactic steering behavior and showed that worms employ the steering mainly for migrating up a thermal gradient to their cultivation temperature [9] (**Fig 1**).

**Fig 1.**
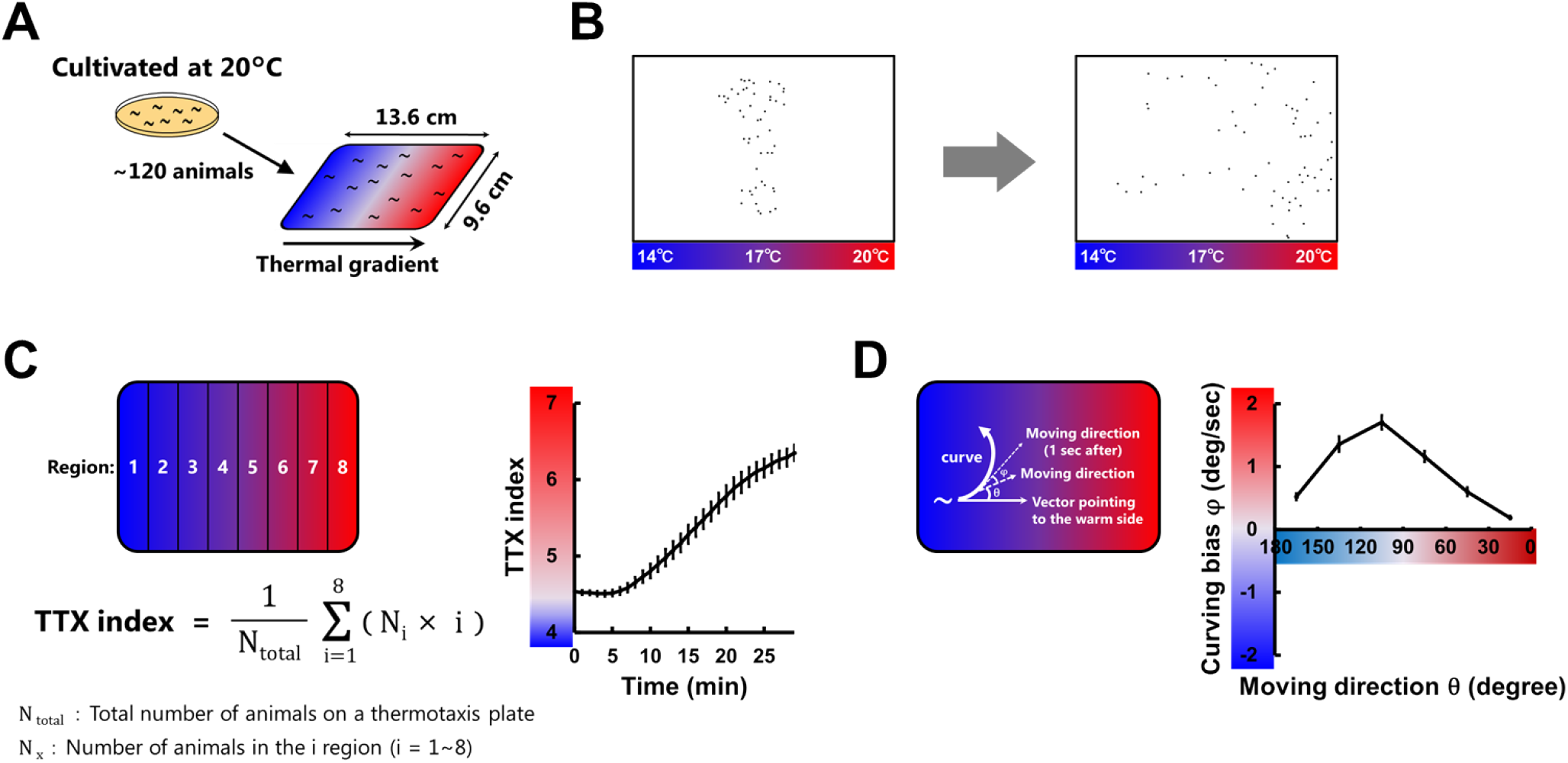
Evaluation of *C. elegans* thermotactic behavior. **(A)** Worms cultivated at 20°C migrate toward the cultivation temperature when placed on a plate with a thermal gradient [7]. **(B)** Representative migration of approximately 120 worms that was recorded by a Multi-Worm Tracker [40]. **(C)** The time course of TTX indices (right panel) calculated using the described equation (left panel) and averaged within the assays (n = 12) [9]. **(D)** Plots of the curving bias φ (right panel) representing the averages as a function of the entry direction θ (left panel) [9]. φ is defined as positive if biased toward higher temperature and negative if biased toward lower temperature. Error bars indicate SEM.

To investigate the circuit basis of steering behavior during thermotaxis, we constructed a neuroanatomically-grounded model with a set of neurons shown to be involved in thermotactic steering behavior [9] (**Fig 2A**). In the model, thermal input sensed by a model worm was converted to the activity of a thermosensory neuron AFD through an empirical response property [18], and inter- and motor neurons were mathematically modeled as passive isopotential nodes with simple first order nonlinear dynamics [19,20] (see Materials and methods). The unknown parameters of the model were evolved using a genetic algorithm so as to reproduce the empirical thermotactic behavior of a population of worms [9] (**Fig 2B**). In the simulation, states of model worms were defined by their position in the assay plate (x, y) and their moving direction relative to the vector pointing to the warm side of the plate (θ). At every step, the model worms were decided whether to perform turns, reversals, or steering behavior according to empirically observed probabilities. When model worms perform turns or reversals, the next θ was provided based on empirical exit directions (Φ). By contrast, when model worms steer, the next θ was provided based on curving angle φ calculated via the model circuit (**Fig 2A and 2B**). A large set of evolutionary searches [21] were performed so that the thermotactic migration and steering behavior of the simulated worms reproduced the empirical data (**Fig 2C**) (see Materials and methods). Across 200 evolutionary searches, we obtained 9 independent parameter sets having a fitness score of at least 0.75 (**Fig 3A and S1A Fig**). Although we did not find prominent common characteristics among the 9 sets of connection weights (**Fig 3B and S1B Fig**), individual models reproduced the time course of TTX index, and the curving bias observed in experiments (**Fig 3C and S1C Fig**). Further, empirical impairments of curving bias in cell-ablated worms [9] were also reproduced in all the 9 models (**S1D Fig**). These results support that our models can serve as platforms to investigate how the neural circuit generates steering behavior during thermotaxis.

**Fig 2.**
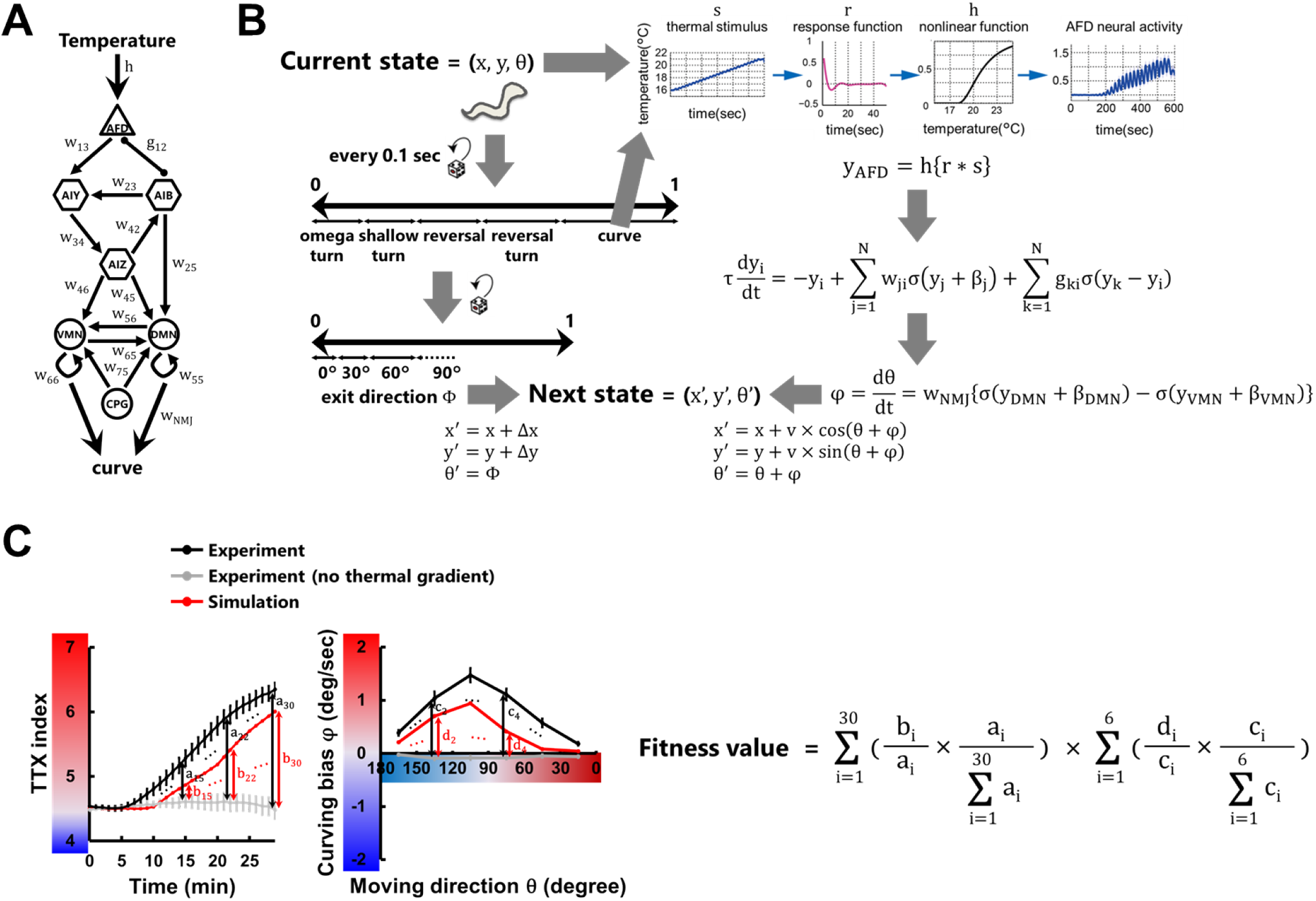
Thermotaxis simulation for building a neuroanatomical model. **(A)** Neuroanatomical model that generates bias in steering behavior during positive thermotaxis [9], including thermosensory neurons (triangles), interneurons (hexagons), and head motor neurons (circles). Black thin arrows indicate chemical synapses, black undirected lines with round endings gap junctions, and gray thick arrows neuromuscular junctions. An oscillatory component CPG is added to generate dorsoventral body undulation of *C. elegans*. **(B)** Schematic structure of the thermotactic behavioral simulation for searching parameters in the model with an evolution algorithm. Worm’s state was defined by its position (x, y) and moving direction (θ) relative to the vector pointing to the warm side. We updated the states of the model worm every 0.1 second in two ways: according to the empirical data for turns and reversals [9] and via the neuroanatomical model for steering behavior. In the former case, frequencies, exit directions (Φ), and displacements (Δx, Δy) during turns and reversals were applied as functions of θ, temperature, and time. In the latter case, activity of AFD was calculated by response property (r and h) [18], activity (y) of interneurons and motor neurons were calculated with simple first order nonlinear dynamics (σ) [39], and curving angle (φ) were calculated proportionally to the difference in activities of dorsal and ventral neck motor neurons (y_DMN_ and y_VMN_). **(C)** Formula for the fitness value in evolutionary searches. TTX index and curving bias from the simulations (red lines) and the experiment (black lines) were subtracted by those from the experiment without thermal gradients (gray lines). The differences were compared (a_i_ versus b_i_, c_i_ versus d_i_), summed up, and multiplied with each other to generate a total fitness value.

**Fig 3.**
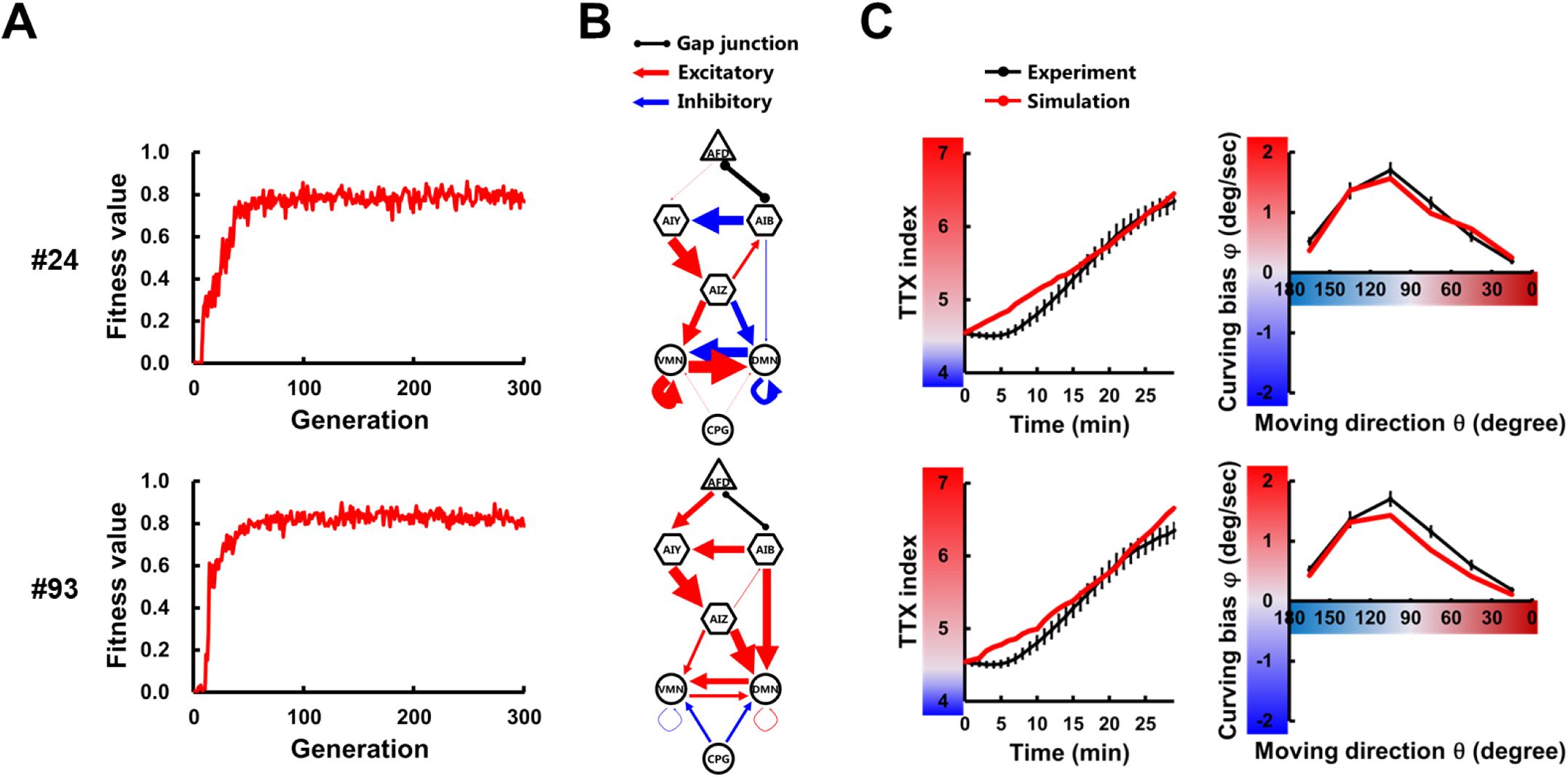
Neuroanatomical models reproduce thermotactic behavior. Through 200 evolutionary searches with 300 generations, 9 independent parameter sets that have fitness scores of at least 0.75 **(A)** were obtained (**S1 Fig**). In the model circuit diagrams **(B)**, the thickness of each connection is represented proportionally to its connection weight. Individual parameter sets were assigned numbers (#) from 1 to 200. With the 2 of 9 models, the time course of TTX index and the curving bias in experiments (black lines) and simulations (red lines) are plotted **(C)**.

### Thermal input on the time scale of forward movement modulates curving rates of locomotion

Since there is no common characteristics among the connection weights of the models (**Fig 3B and S1B Fig**), we first assessed whether there exists common profiles of steering behavior under simplified simulation settings. Temperature of assay plates was set as constant, and model worms were set not to perform turns and reversals. As shown in **Fig 4A**, the curving rate of model worm’s trajectory was smaller as the temperature of plates was higher. This negative relation between absolute temperature and curving rates |φ| was observed in all the 9 parameter sets, though the magnitudes of curving rates were diverse (#93 and #156 versus others) and some traces were non-monotonic (#48 and #156) (**S2 Fig**).

**Fig 4.**
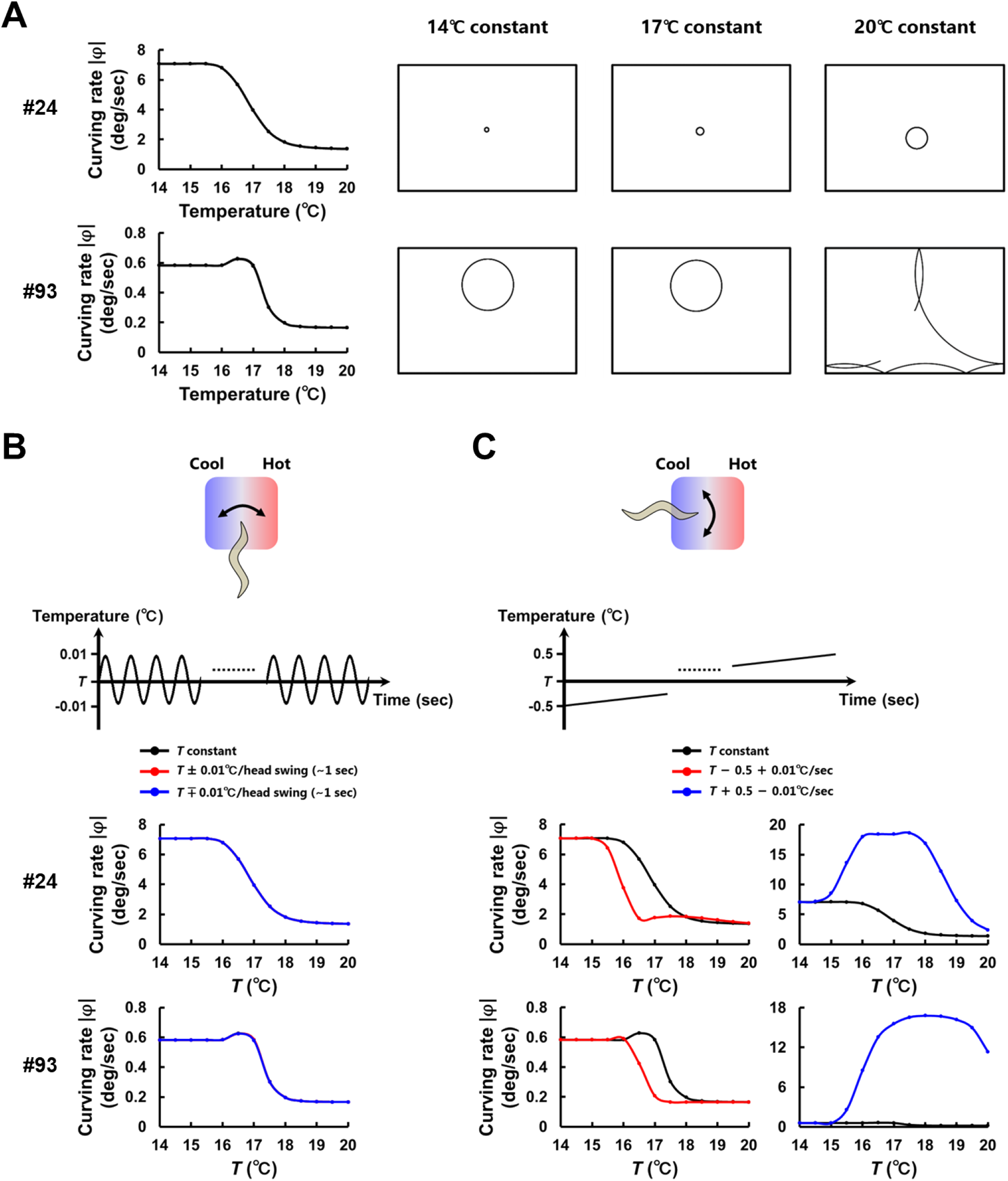
Temperature change on the time scale of forward movement modulates curving rates of locomotion. **(A)** Trajectory of simulated worms on the constant temperatures (14–20°C). The curving rates of trajectories are calculated and plotted against temperatures (left panels). **(B)** Curving rates under temperature change on the time scale of head swings (upper panels). Worms are assumed to be moving perpendicularly to a thermal gradient with their dorsal side heading toward warmer side (red lines) or colder side (blue lines). Curving rates under these conditions were compared with those at the constant temperature (black lines). In the lower panels, red, blue, and black lines are overlapping. **(C)** Curving rates under temperature change on the time scale of forward movement (upper panel). Worms are assumed to be moving straight up a thermal gradient (red lines) or down a thermal gradient (blue lines). Curving rates under these conditions were compared with those at the constant temperature (black lines). The results with representative parameter sets (#24 and #93) are shown.

To examine whether curving rates are also affected by derivatives of temperature, the model worms were next exposed to temperature changes that mimic those sensed by the worms on a thermal gradient. One type of temperature change is at the scale of head swings of worms (**Fig 4B**). Due to their dorsoventral movement, worms sense sinusoidal changes in temperature with a maximum amplitude of 0.01°C on a linear thermal gradient of 0.5°C/cm [26]. Although the sensory neuron AFD in the models responded to the sinusoidal thermal input (**S3A Fig**), curving rates of the model worms were not different from those at the constant temperatures (**Fig 4A and 4B**). The other type of temperature change is at the scale of forward movement of worms (**Fig 4C**). When moving up (or down) a thermal gradient of 0.5°C/cm, worms sense a temperature increment (or decrement) of approximately 0.01°C/sec. We found that, within the range 16–18°C, temperature increments decreased curving rates of animals compared with those at a constant temperature, whereas temperature decrements increased curving rates within the range 15–20°C. AFD showed larger responses in these linear thermal inputs (**S3B Fig**) than in the sinusoidal thermal inputs (**S3A Fig**). This analysis suggests that thermal input from the worm’s forward movement, but not from the worm’s head swings, modulate the curving rate of locomotion, thereby steering to preferred temperature.

### Thermotactic behavior can be generated without directed biases in curving rates

The temperature changes employed in **Fig 4B and 4C** correspond to those sensed by worms moving perpendicularly or in parallel to a thermal gradient; the moving direction θ (**Fig 1D and 2B**) is equal to 90° or 0°/180°, respectively. We further examined curving profiles under other θ and represented curving rates |φ| as functions of θ and temperature *T* (**Fig 5A and S4 Fig**). The positive relation between θ and |φ| was observed in all the parameter sets, though some surfaces were non-monotonic (#48) or steep (#93, #140, and #156).

**Fig 5.**
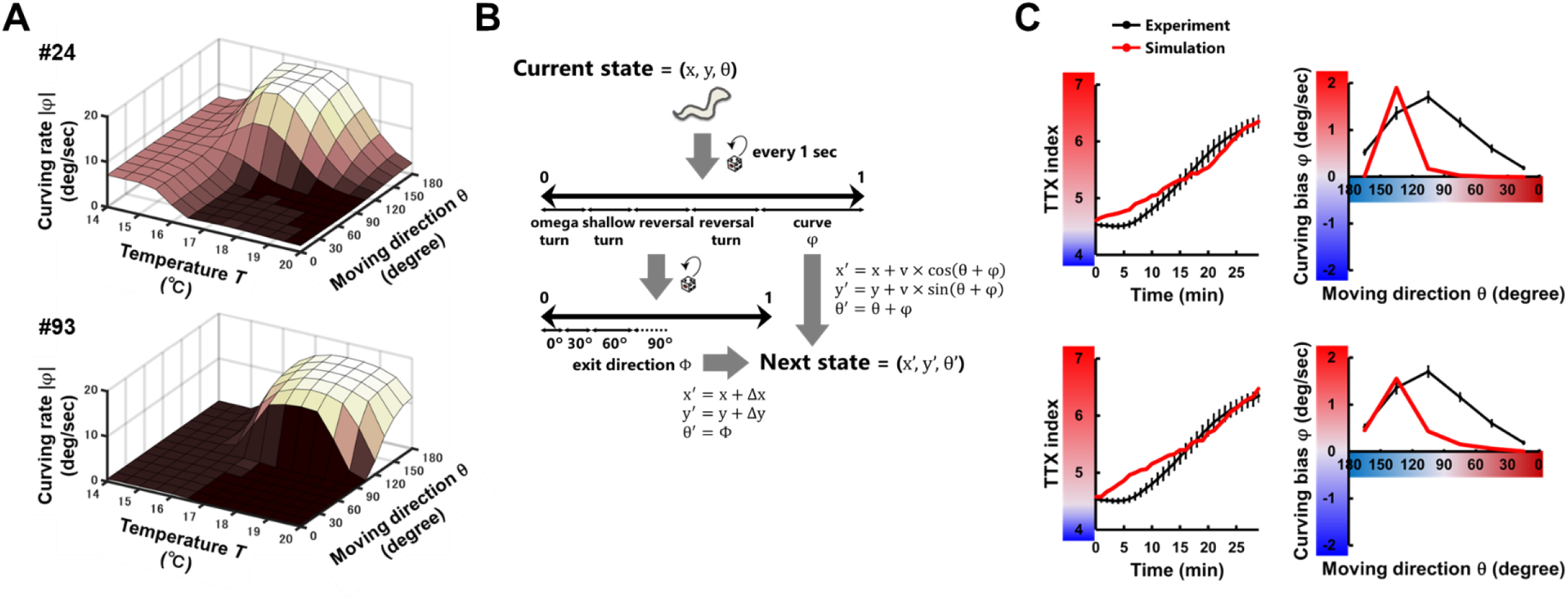
Profiles of curving rates against temperature change reproduce thermotactic behavior. **(A)** Curving rates |φ| were calculated and plotted against temperature *T* and moving direction θ. **(B)** Schematic structure of the thermotaxis behavior simulation based on the profiles of curving rates shown in **A**. We updated the states of the worm every 1 second according to the empirical data for turns and reversals [9] and via the profile of curving rates, in which curving angle (φ) were calculated by multiplying random signs. **(C)** The time course of TTX index (left column) and the curving bias (right column) in experiments (black lines) and simulations (red lines) are plotted. The results with representative parameter sets (#24 and #93) are shown.

To assess whether the curving profiles in **Fig 5A and S4 Fig** can generate thermotactic migration and curving biases as observed in experiments, we conducted another Monte Carlo simulations of population thermotaxis in which curving rates of worms are decided not via the neuroanatomical model (**Fig 1D**) but based on the curving profiles (**Fig 5B**). When the model worm performs steering behavior, the amplitude of curving angle (|φ|) is decided based on the profiles in **Fig 5B**, and the direction of steering (that is whether to steer higher or lower temperature direction) is randomly determined. Thus, the simulated worms lack the opportunity to bias steering consistently toward warmer or cooler directions. Nevertheless, we found that the simulations reproduced thermotactic migrations toward warmer direction and generated curving bias toward higher temperature (**Fig 5C**). This analysis demonstrates that modulation of curving rate upon thermal input along forward movement (**Fig 4C**) can generate thermotactic migration.

### Activity of a thermosensory neuron decides curving rates of locomotion

The observation that curving rate |φ| is dependent on temperature *T* and moving direction θ (**Fig 5A**) implies the dependence of |φ| on activity of a thermosensory neuron AFD (y_AFD_), since AFD encodes both absolute temperature and the differential of temperature [9,18,27]. To investigate this relationship, we examined |φ| of model worms under simulations in which y_AFD_ is fixed at constant value, within the range in which model worms experience during the simulations of population thermotaxis (**Fig 3C**). As shown in **Fig 6**, |φ| was smaller as y_AFD_ was higher. This negative relationship was observed in all the 9 parameter sets, though some traces were non-monotonic (#48 and #94) (**S5A Fig**). Notably, in 5 of 9 parameter sets (#17, #22, #33, #94 and #140), |φ| took the minimum values at specific y_AFD_ values; |φ| was larger as y_AFD_ was higher than these values. The conversion from negative to positive relation was accompanied by conversion of running direction from clockwise to counterclockwise, or vice versa (**S5B Fig**). Overall, y_AFD_ experienced by model worms during the thermotaxis simulations covered the range in which |φ| showed dynamic decrease upon increase of y_AFD_ and the values in which |φ| took the minimum (**S5A Fig**). These results suggest that thermal input sensed by AFD is efficiently transformed to regulate curving rates, generating the curving profiles shown in **Fig 5A** and leading to thermotactic behavior (**Fig 5C**).

**Fig 6.**
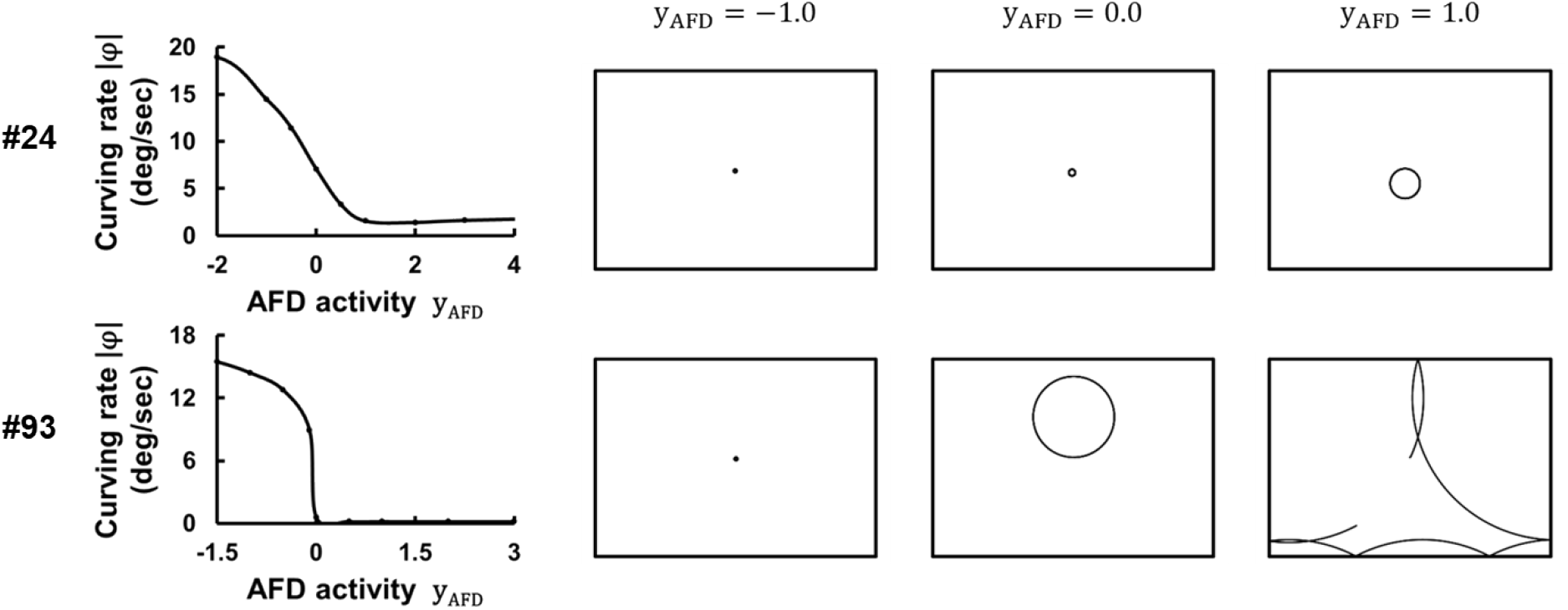
Activity of a thermosensory neuron AFD modulates curving rates of locomotion. Trajectory of simulated worms on the fixed AFD activity (y_AFD_). The curving rates |φ| of trajectories were calculated and plotted against y_AFD_.

### Thermal input is transmitted from sensory to motor neurons on a longer time scale than head swings

The next question to ask is how thermal input sensed by AFD is transmitted to inter- and motor neurons (**Fig 2A**) to generate steering behavior. Correlation analysis and information-theoretic analysis (see Materials and methods) revealed that the dynamics of AFD is naively transmitted to amphid interneurons AIB, AIY, and AIZ (**Fig 7A and 7B**), in which transmission valences are consistent with connection valences (**Fig 3B**). By contrast, since oscillatory input from a central pattern generator (CPG) evokes oscillatory activity in dorsal/ventral motor neurons (y_DMN_ and y_VMN_) (**Fig 2A**) and thus curving rate (|φ|) exhibits oscillatory dynamics (**Fig 7A**), we did not observe a significant relationship (linear or non-linear) between y_AFD_ and y_DMN_/y_VMN_/|φ| (**Fig 7B**).

**Fig 7.**
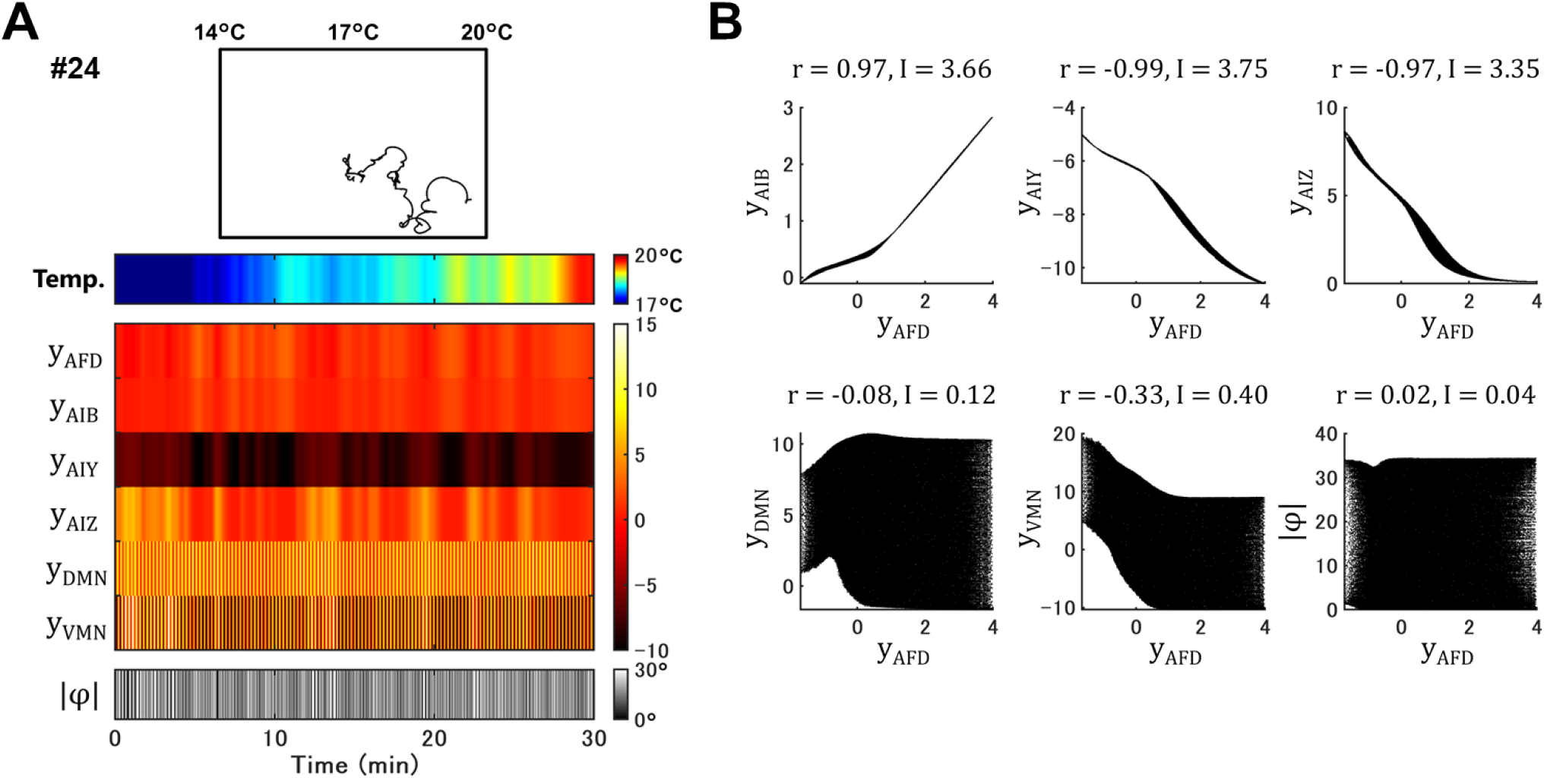
Activity of a thermosensory neuron and interneurons shows strong correlations. **(A)** Representative trajectory of a simulated worm (upper panel) through the neuroanatomical model with a representative parameter set (#24). The time courses of temperature sensed by the worm, activity of individual neurons, and curving rate |φ| are represented in color scales (lower panels). **(B)** Scatter plots of AFD activity (y_AFD_) versus activity of other neurons or curving rate |φ| of 100 simulated worms. Correlation coefficients (r) and mutual information (I) between them were calculated.

To extract the components from y_DMN_/y_VMN_/|φ| in which the dynamics of y_AFD_ are embedded, we performed singular spectrum analysis (SSA) (see Materials and methods). SSA decomposes time-series data into the left eigenvector (U) that corresponds to eigen-time-series (singular spectrum) and the right eigenvector (V) that represents the magnitude of each of the eigen-time-series (**Fig 8A**). SSA on the y_DMN_/y_VMN_/|φ| shown in **Fig 7A** revealed that variables in V whose corresponding eigen-time-series in U is constant within a 4-sec time window exhibited the strongest correlation and mutual information with y_AFD_ (**Fig 8B and 8C**). This tendency was observed in all the 9 parameter sets (**Fig 8D**), indicating that the transmission of activity from sensory to motor neurons is evident in the longer time scale than sinusoidal head swings of worms.

**Fig 8.**
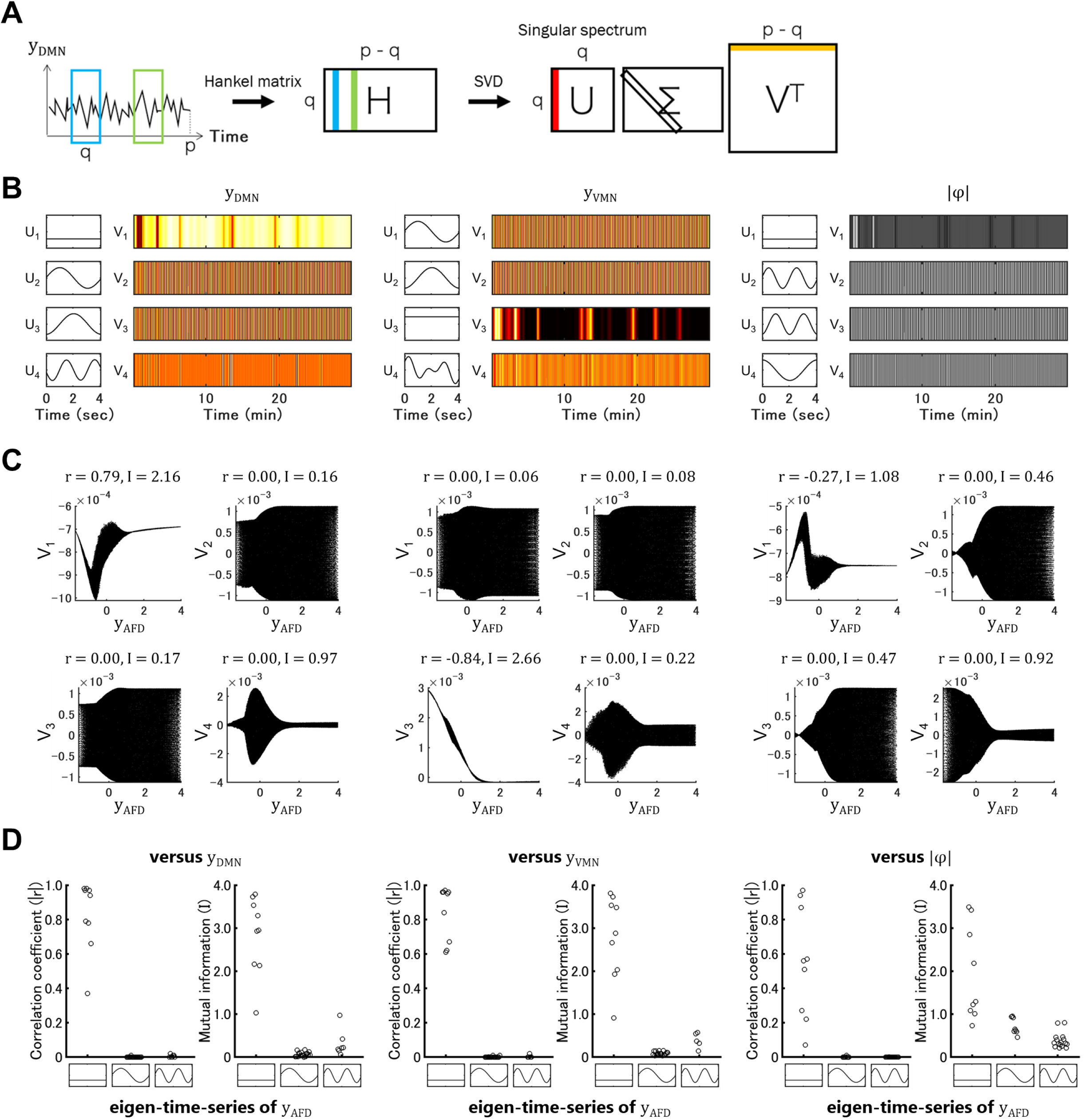
Dynamics of constant singular spectrum of motor neuron activity show stronger correlation with a thermosensory neuron activity. **(A)** Singular spectrum analysis (SSA) is an algorithm to analyze time-series data. A time series of neural activity data y(t) (t = 1~p) is stacked into a Hankel matrix (H) with a sliding time window of q. The singular value decomposition (SVD) of H yields a hierarchy of eigen time series that represent the singular spectrums (U) and their magnitude time series (V). **(B)** SSA on activity of dorsal/ventral motor neurons (y_DMN_ and y_VMN_) and curving rate (|φ|) of a simulated worm in **Fig 7A**. The first four eigen-time-series U and their magnitude V are shown. **(C)** Scatter plots of AFD activity (y_AFD_) versus each variable in V of 100 simulated worms. Correlation coefficients (r) and mutual information (I) between them were calculated. **(D)** In all the 9 models, r and I between y_AFD_ and magnitudes of first four eigen-time-series of y_DMN_/y_VMN_/|φ| are plotted.

## Discussion

In this study, we constructed a neuroanatomically-grounded model using the neurons that have been shown to mediate thermotactic steering behavior in *C. elegans* (**Fig 1–3**) and demonstrated that thermal input sensed not through head swings but through forward movement can adjust the worm’s moving direction and implement navigation to its preferred temperature (**Fig 4–6**). Singular spectrum analysis (SSA) revealed that the information transmission from sensory to motor neurons occurred over a longer time scale than the worm’s dorsoventral head swing (**Fig 8**).

The previous studies on *C. elegans* steering behavior have proposed that steering behavior is based on adjustment of head swing amplitude upon dorsoventral sinusoidal motion, leading to direct orientation to a destination [10,11], and thus is categorized as klinotaxis [1,28]. This hypothesis is supported by imaging [12–14] and optogenetic experiments [13,15,16]. Also in modeling studies, neuroanatomical models which were built to reproduce empirical chemotactic steering behavior show that oscillatory activity of motor neurons are modified upon the application of favorable/unfavorable gustatory signals [19,29]. By contrast, in this study, thermotactic steering behavior was based on adjustment of curving rate upon forward locomotion (**Fig 4B and 4C**), leading to indirect orientation to destination temperature (**Fig 5B and 5C**), and thus can be categorized as klinokinesis [1,28]. Thermotactic steering as indirect orientation is convincing when we consider subtle and variable thermal environment where *C. elegans* inhabits [17]. On a linear thermal gradient of 0.5°C/cm, thermal input sensed through head swings varies ±0.01°C within a couple of seconds [26], while thermal input sensed through forward movement is more persistent and monotonous. The temperature change itself on these two situations is at the same rate, 0.01°C/sec (**Fig 4A**).

The klinokinetic regulation of thermotactic steering behavior is potentially verified in assays in which superposed thermal variation is applied temporally to the entire surface of an agar plate, and behavior of an animal is monitored simultaneously. By applying sinusoidal or monotonous thermal input and comparing curving rates of animals under these two conditions, we could speculate which type of thermal input is more directly transformed to steering behavior. However, the application of such sinusoidal temperature change with small amplitude and high frequency is difficult to achieve since the thermal conductivity of agar is not so large.

Unlike other decomposition methods, such as short-time Fourier transform (STF) or continuous wavelet transform (CWT), SSA is a nonparametric decomposition of time series data [30,31] which does not assume temporal stationarity and spatial consistency within a multivariate system [32]. SSA decomposes time series into a sum of singular spectrums (U), which represent eigen-time-series, and time series vectors (V), which represent the magnitude of each of the singular spectrum (**Fig 8A**). SSA was originally employed for denoising and extracting essential dynamics from measured data, especially geophysical time series [33–35]. A recent study utilized SSA for linear representation of nonlinear dynamics in chaotic systems [36]. Also, by investigating singular spectrums, SSA can be applied for change-point detection in time series [37]. In this study, we decomposed activity of the head motor neurons DMN/VMN and curving rate |φ| by SSA and found that the magnitude of not oscillatory but constant spectrums exhibit the most evident relations with activity of the sensory neuron AFD (**Fig 8C and 8D**). Since DMN and VMN receive out of phase sinusoidal input from a pattern generator CPG (**Fig 2A**) and generate sinusoidal oscillation of |φ| (**Fig 7A**), their singular spectrums were mostly sinusoidal (**Fig 8B**), which can be also extracted by STF or CWT. However, for example, the fourth spectrum of VMN in **Fig 8B** cannot be represented in these two methods. The dynamics that corresponds to constant spectrums cannot be naturally extracted by STF and CWT. Further, in real nervous system, a wide variety of eigen-time-series is shown to underly (i.e. temporal non-stationarity and spatial inconsistency) [38]. Therefore, SSA would be a powerful method for decomposing and understanding neural dynamics.

## Materials and Methods

### Neuroanatomical modeling

Mathematical model of a neural circuit for regulating steering behavior (**Fig 2A**) was constructed based on previous studies [19,20]. Thermosensory neuron AFD was modeled as a node with response property that was identified previously [18]. Interneurons and motor neurons were modeled as passive isopotential nodes with simple first order nonlinear dynamics [39]. The dorsal and ventral neck motor neurons receive out of phase input from an oscillatory component CPG which models dorsoventral body undulation of *C. elegans*. Chemical synapses were modeled as sigmoidal functions of presynaptic voltage, and gap junctions were modeled as non-rectifying conductances between two neurons. The curving angle was calculated proportionally to the difference in activities of dorsal and ventral neck motor neurons.

### Thermotaxis simulation

Thermotaxis behavior was simulated with the neuroanatomical model (**Fig 2A**) and behavioral data [9] recorded by a Multi-Worm Tracker [40,41]. For each simulation, 100 model worms were run sequentially. Model worms were considered as dimensionless points in a 13.6 cm (x axis) × 9.6 cm (y axis) plate, with a linear thermal gradient from 14 to 20 °C along the x axis. Model worms started from the center of a plate, while y coordinates and initial directions were randomized. For every second, individual model worms were decided whether to undergo an omega turn, a shallow turn, a reversal, a reversal turn, or a curve (**Fig 2B**). Event probabilities of each behavioral component were defined according to the experimental data of turning frequencies. When model worms were decided to do any turns, the next moving directions (θ) were defined according to the experimental data of exit directions (Φ). The next positions (x, y) were defined together with the experimental data of the displacements (Δx, Δy) during the individual turns. Every experimental data was applied as a function of moving direction θ. Besides, different data set were applied depending on whether model worms were on the fraction 1–2, the fraction 3–6, or the fraction 7–8 of a thermotaxis plate (**Fig 1C**), and 0–10 min, 10–20 min, or 20–30 min of a simulation. When model worms were decided to do a curve, the next moving directions (θ) were defined according to the curving angle φ, calculated through the neuroanatomical model. The next positions (x, y) were defined together with the moving speed 0.2 mm/sec. If a model worm reaches the plate border, it was set to do specular reflection.

### Evolution algorithm

Parameters of the neuroanatomical model were optimized by applying a genetic algorithm [21]. We optimized the following 16 parameters: weights of chemical synapses (w), weights of gap junctions (g), weights of neuromuscular junction (w_NMJ_), terms that determine response property (h) of a sensory neuron, and bias terms (β) that shift sensitivity range of inter- and motor neurons [42] (**Fig 2B**). The optimization algorithm was run for populations of 96 independent parameter sets. Each time the algorithm was run, individuals were initialized by random selection. Populations were evolved for 300 generations. At the end of a run, the parameters of the best performing individual were stored for later analysis. The algorithm was run 200 times, yielding 200 distinct model networks. Fitness value of the model was evaluated based on population thermotaxis simulation (**Fig 2B**), in which activity of AFD sensory neuron was calculated from the time series of experienced temperature (s) of the simulated animal, activity of downstream neurons (y) were then calculated, and curving angle φ was calculated, determining the next moving directions (θ). TTX index from the simulation was compared with the empirical index (**Fig 1C**) in every minute, and the difference between them was summed up within 1–30 min. In a similar way, curving bias from the simulation was compared with the empirical bias (**Fig 1D**) along every 30 degree of θ, and the difference between them was summed up within 0–180°. These two values were normalized with the summation of the difference between the empirical TTX indices (or curving biases) with those in no thermal gradient conditions and then multiplied with each other to generate a total fitness value (**Fig 2C**).

### Decomposition analysis

Time-series of neural activity data were analyzed by singular spectrum analysis (SSA) [30,31] (**Fig 8A**). In SSA, we construct a Hankel matrix (H) from 1-d time-series data and decompose the matrix with singular value decomposition (SVD). The left eigenvector (U) represents the feature vector of time-series, namely singular spectrum, and the right eigenvector (V) represents the magnitude of each of the singular spectrum.

### Information-theoretic analysis

Time-series of multiple neural activity data (**Fig 7B**) or magnitudes of singular spectrum (**Fig 8C**) were analyzed by calculating mutual information (I) [43]. Calculation of mutual information requires probability distributions of the time-series data, and we generated them as previously described [29]. The discrete probability distributions were estimated over a fine grid of 50 bins and by a kernel density estimation technique known as average shifted histograms [44], with 12 shifts along each dimension. Mutual information is a measure of the dependence between two variables [43] and quantifies the amount by which a measurement on one of the variables reduces our uncertainty about the other. Mutual information can measure a non-linear relation between variables, whereas Pearson’s correlation coefficient (r) measures a linear relation.

### Quantification

Experimental data are expressed as mean ± SEM. Simulation data are expressed as mean.

## Acknowledgments

We would like to thank Ikue Mori for supervising experimental data acquisition; Erick Olivares and Randall D. Beer for valuable comments on modeling analyses; and Jun Kitazono and Masafumi Oizumi for helpful discussions. M.I. was supported by Grant-in-Aid for Scientific Research (KAKENHI) 16J05770. This work was supported by NSF grant IIS-1845322 (to E.J.I.).

## Supporting Information

**S1 Fig.**
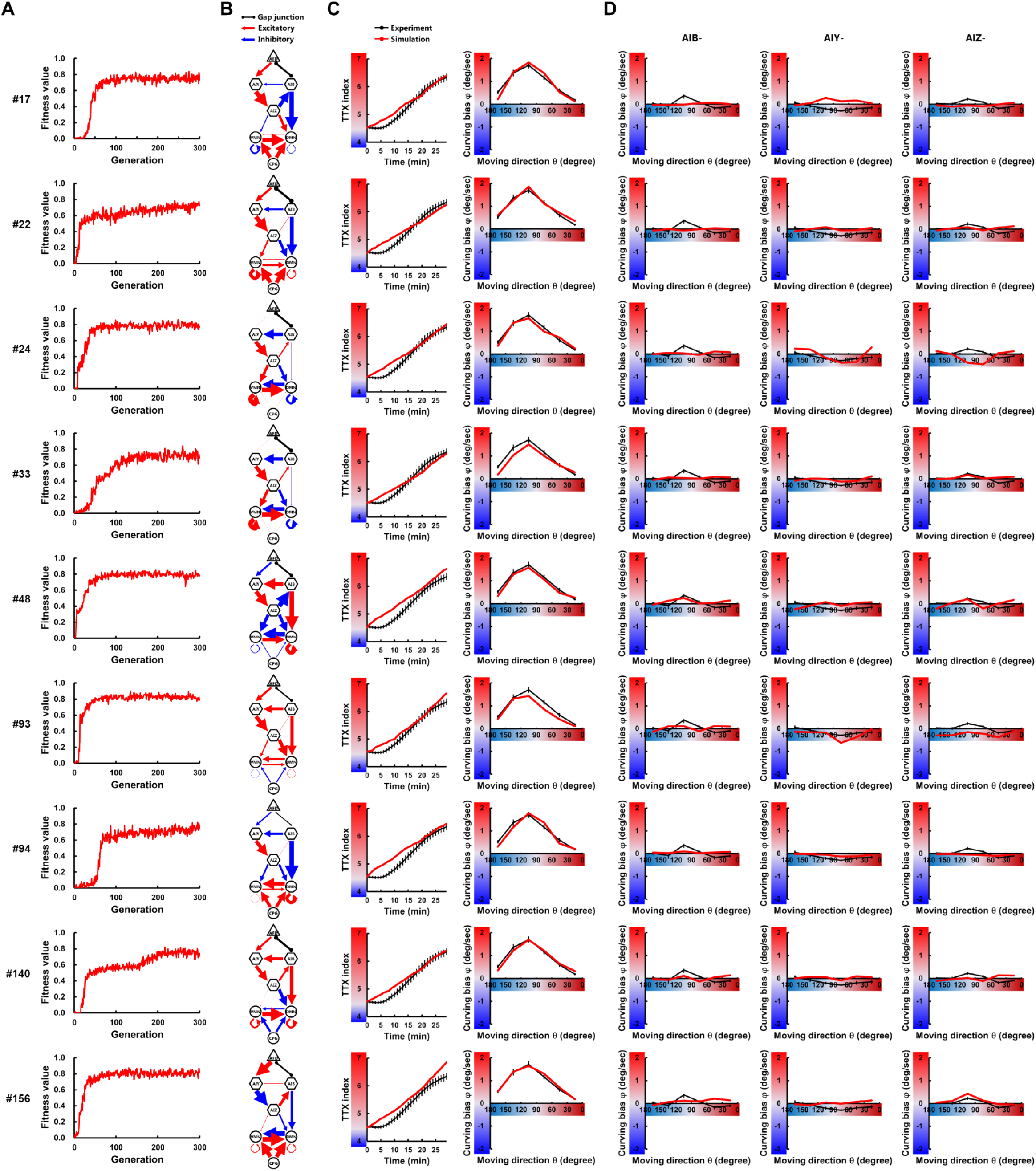
Thermotactic behavior is reproduced by a variety of neuroanatomical models. Through 200 evolutionary searches with 300 generations, 9 independent parameter sets that have fitness scores of at least 0.75 **(A)** were obtained. In the model circuit diagrams **(B)**, the thickness of each connection is represented proportionally to its connection weight. Individual parameter sets were assigned numbers (#) from 1 to 200. The time course of TTX index and the curving bias in experiments (black lines) and simulations (red lines) are plotted **(C)**. All the 9 models reproduced empirical impairments of curving bias upon ablating AIB, AIY, and AIZ **(D)**.

**S2 Fig.**
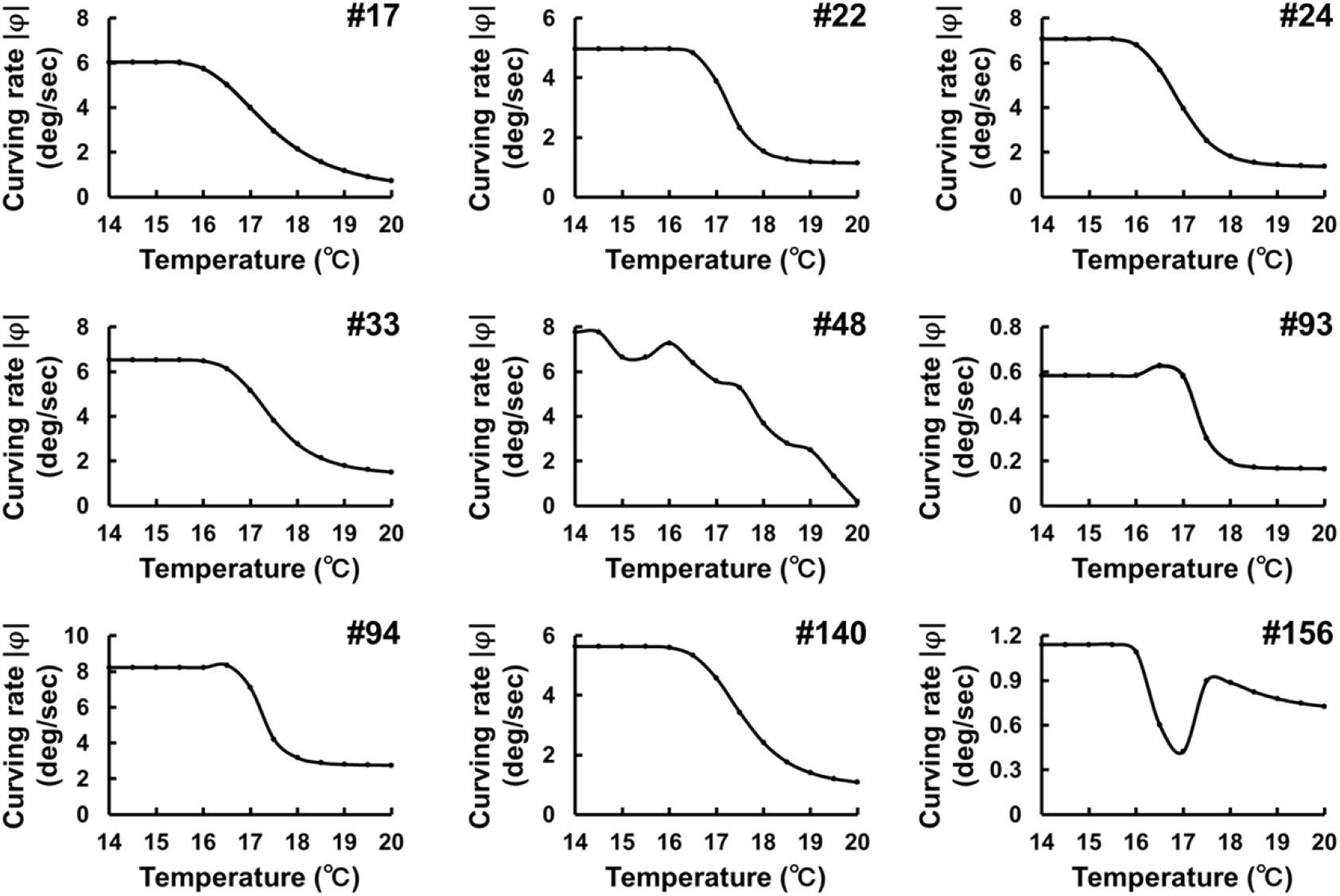
Negative relation between absolute temperature and curving rates. Trajectory of simulated worms on the constant temperatures (14–20°C). In all the 9 models, the curving rates of trajectories were calculated and plotted against temperatures (left panels).

**S3 Fig.**
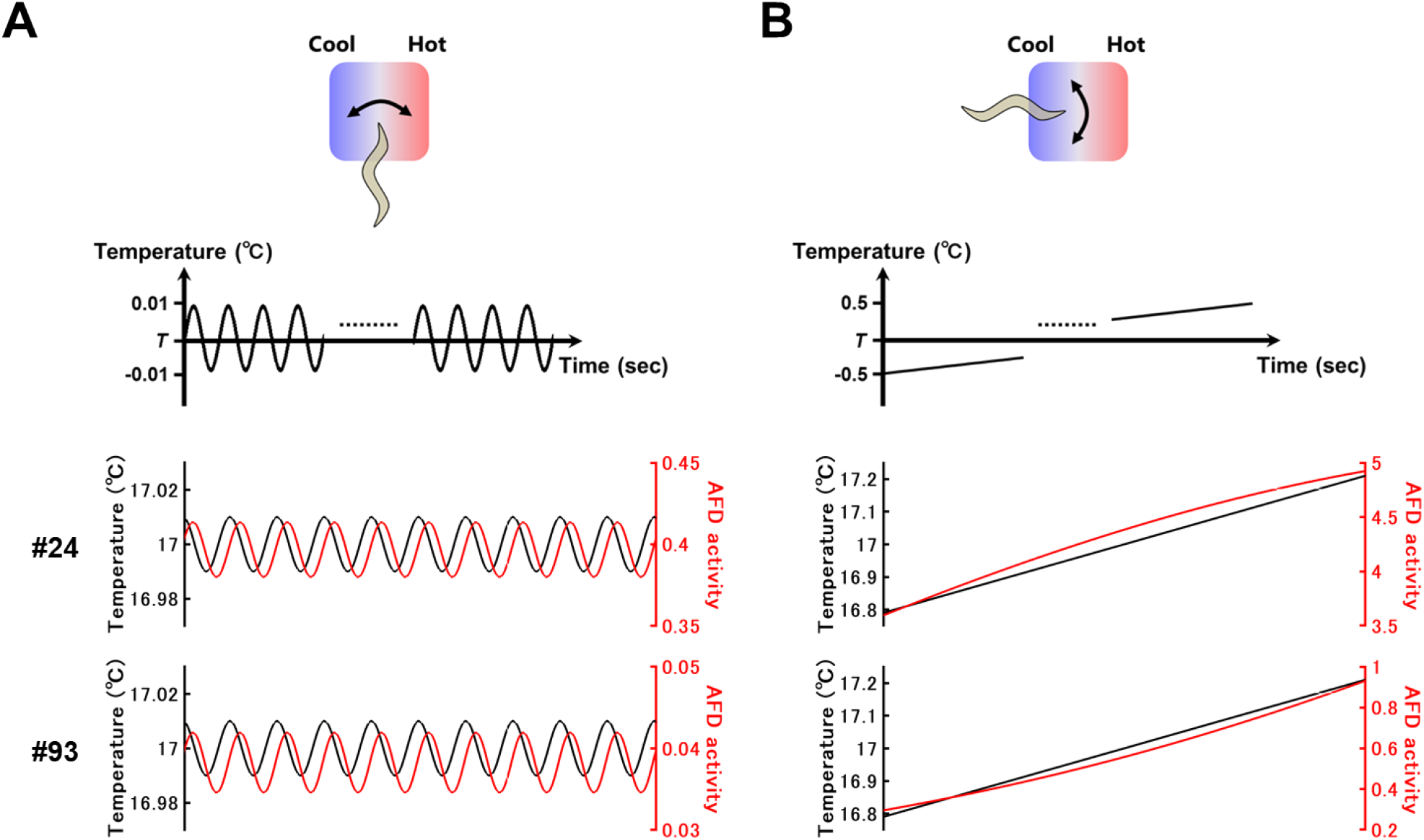
AFD responds to temperature change on the time scale of head swings and forward movement. **(A)** Activity of AFD under temperature changes on the time scale of head swings. Worms are assumed to be moving perpendicularly to a thermal gradient with their dorsal side heading toward warmer side. **(B)** Activity of AFD under temperature changes on the time scale of forward movement. Worms are assumed to be moving straight up a thermal gradient. The results with representative parameter sets (#24 and #93) are shown.

**S4 Fig.**
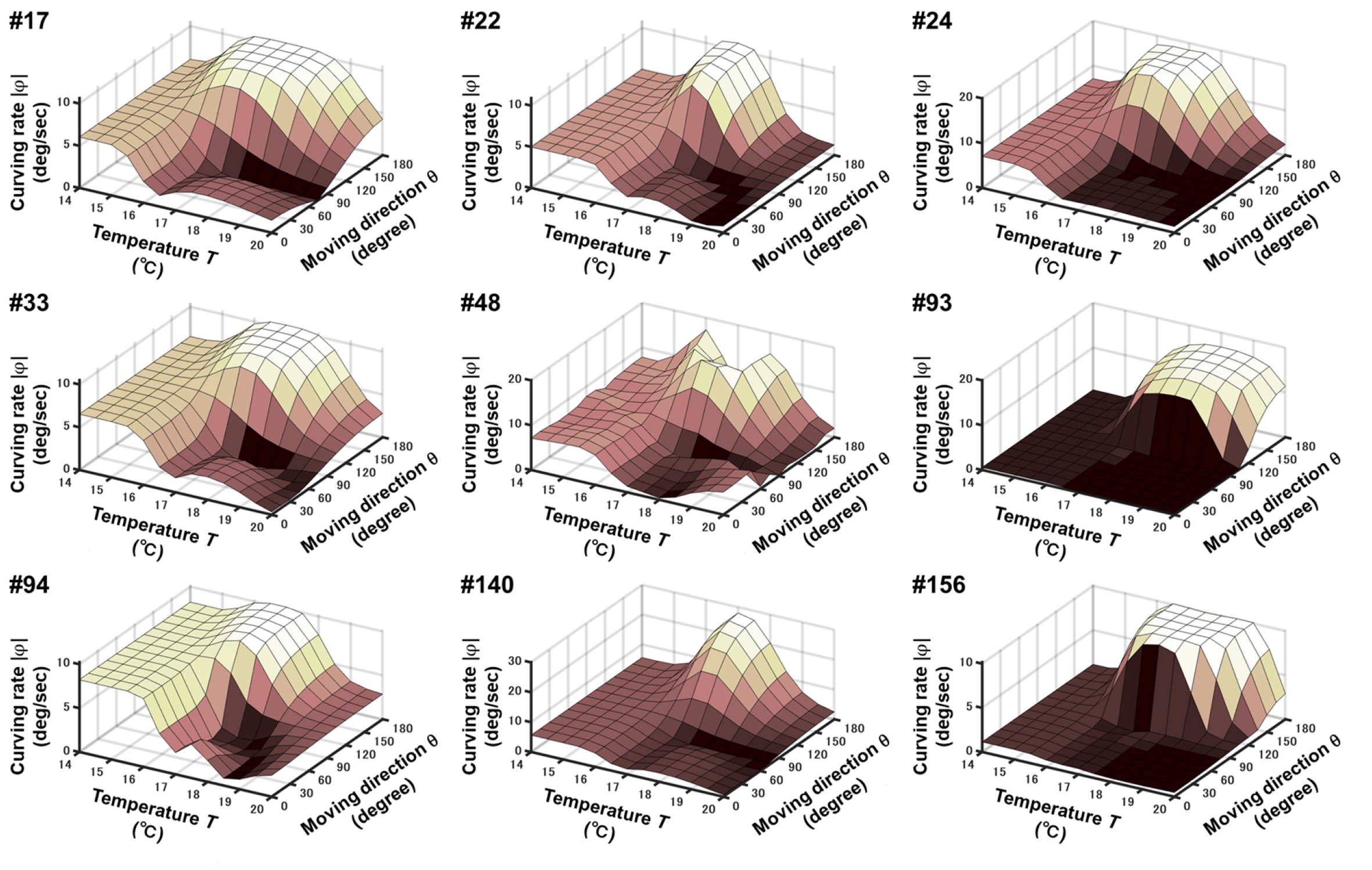
Profiles of curving rates against absolute temperature and derivative of temperature. In all the 9 models, the curving rates |φ| were calculated and plotted against temperature *T* and moving direction θ.

**S5 Fig.**
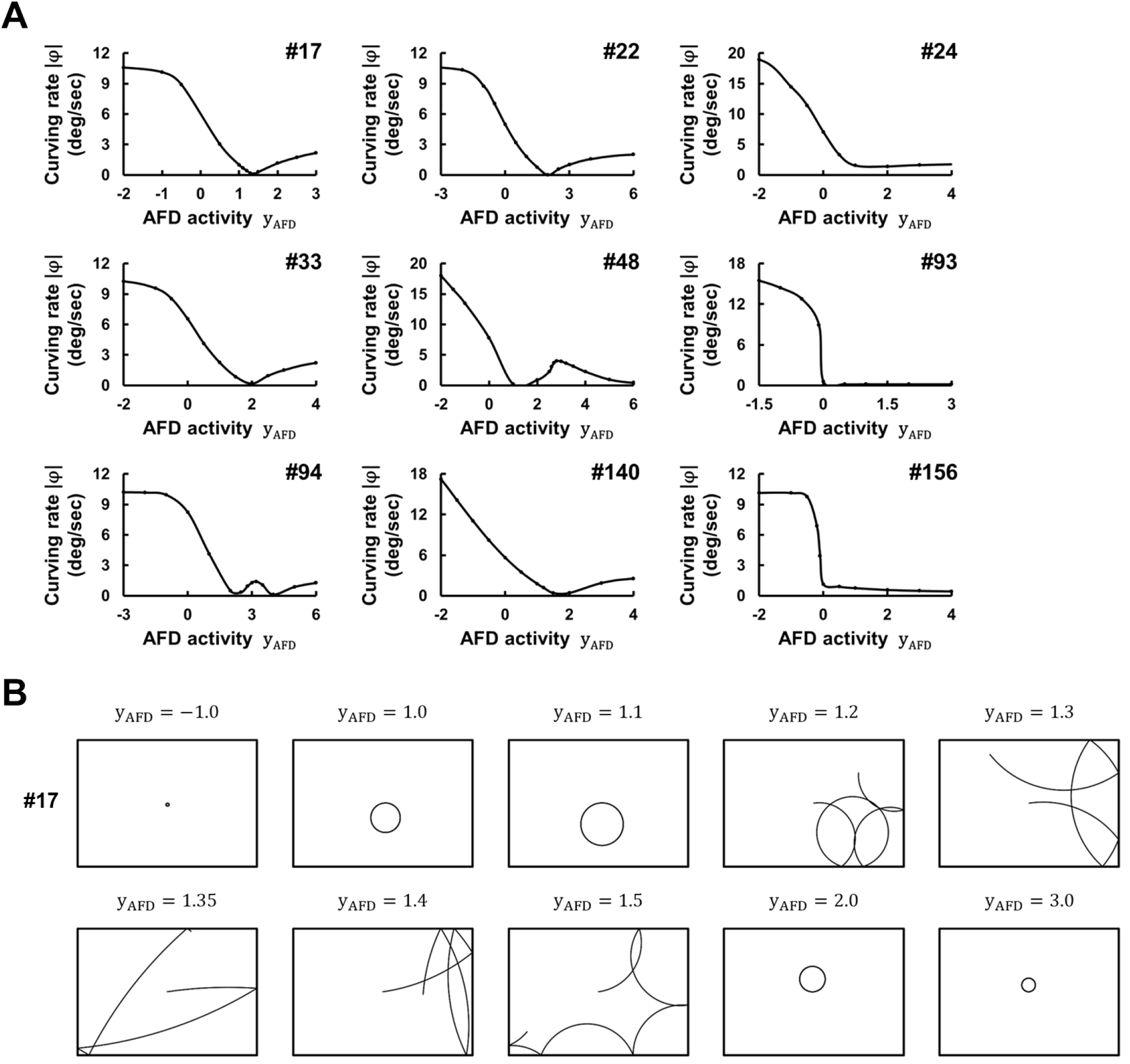
Activity of a thermosensory neuron AFD modulates curving rates and circulating direction of locomotion. **(A)** In all the 9 models, the curving rates |φ| of trajectories were calculated and plotted against AFD activity (y_AFD_). **(B)** Trajectories of simulated worms with representative parameter set (#17) under the fixations of y_AFD_.

